# Specific viral RNA drives the SARS CoV-2 nucleocapsid to phase separate

**DOI:** 10.1101/2020.06.11.147199

**Authors:** Christiane Iserman, Christine Roden, Mark Boerneke, Rachel Sealfon, Grace McLaughlin, Irwin Jungreis, Chris Park, Avinash Boppana, Ethan Fritch, Yixuan J. Hou, Chandra Theesfeld, Olga G Troyanskaya, Ralph S. Baric, Timothy P. Sheahan, Kevin Weeks, Amy S. Gladfelter

## Abstract

A mechanistic understanding of the SARS-CoV-2 viral replication cycle is essential to develop new therapies for the COVID-19 global health crisis. In this study, we show that the SARS-CoV-2 nucleocapsid protein (N-protein) undergoes liquid-liquid phase separation (LLPS) with the viral genome, and propose a model of viral packaging through LLPS. N-protein condenses with specific RNA sequences in the first 1000 nts (5’-End) under physiological conditions and is enhanced at human upper airway temperatures. N-protein condensates exclude non-packaged RNA sequences. We comprehensively map sites bound by N-protein in the 5’-End and find preferences for single-stranded RNA flanked by stable structured elements. Liquid-like N-protein condensates form in mammalian cells in a concentration-dependent manner and can be altered by small molecules. Condensation of N-protein is sequence and structure specific, sensitive to human body temperature, and manipulatable with small molecules thus presenting screenable processes for identifying antiviral compounds effective against SARS-CoV-2.

## Introduction

The outbreak of COVID-19, caused by the severe acute respiratory syndrome-related coronavirus SARS-CoV-2, is a global public health crisis. Coronaviruses, including SARS-CoV-2, are RNA viruses with ∼30 kb genomes that are replicated and packaged in host cells. Packaging is thought to be highly specific for the complete viral genome (gRNA), and excludes host RNA and abundant virus-produced subgenomic RNAs (*1*). Viral replication and gRNA packaging depends on the nucleocapsid protein (N-protein) (*2, 3*). The N-protein has two RNA-binding domains, forms multimers (*4*) and is predicted to contain intrinsically disordered regions **(Figure 1A)**. N-protein thus has hallmarks of proteins that undergo liquid-liquid phase separation (LLPS), a process which may provide selectivity and efficiency to viral replication and packaging.

**Figure 1:**
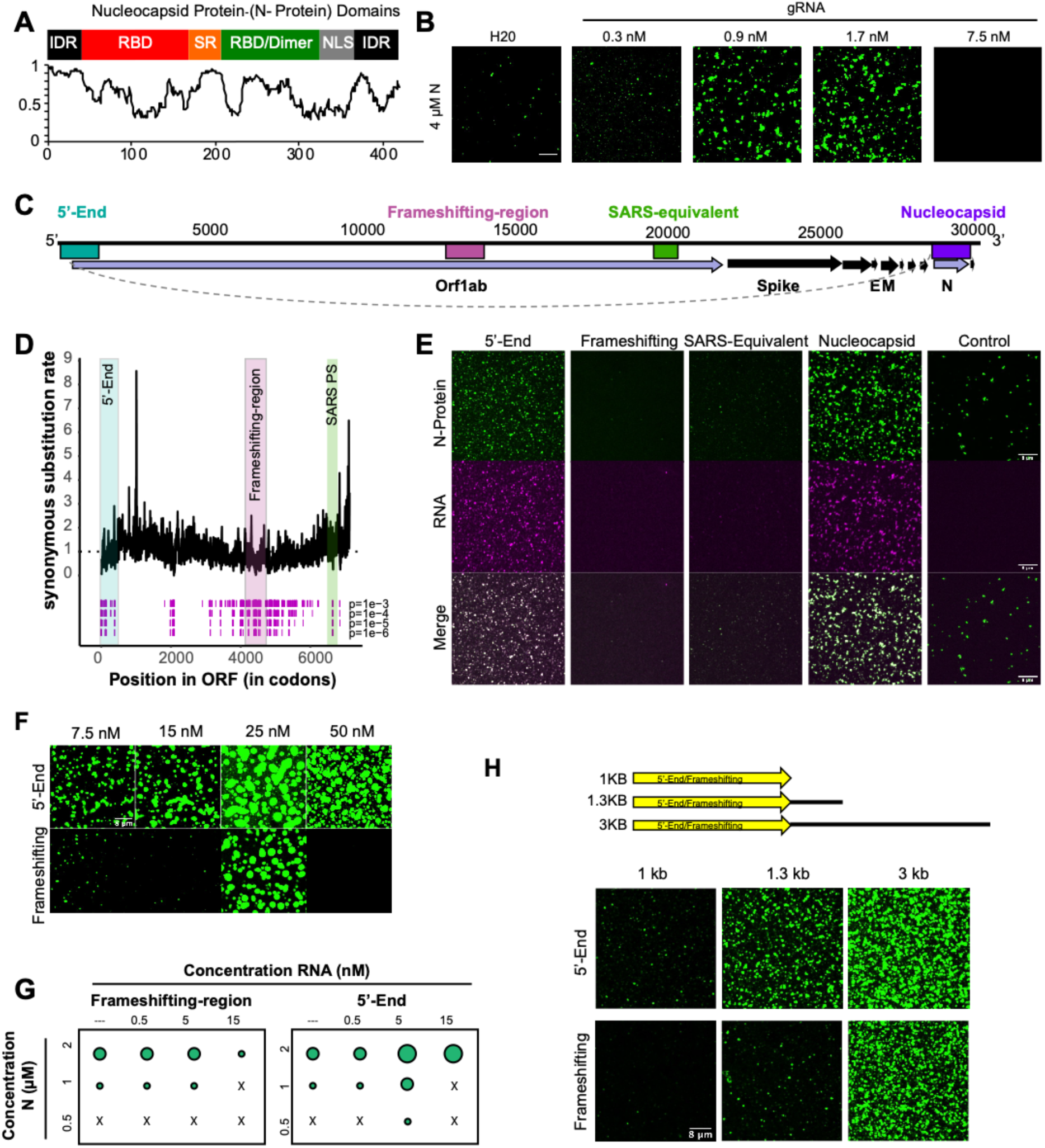
N-protein and RNA mediated phase separation: **A)** Top: domain structure of N-protein. Bottom: disorder plot (y-axis) of N-protein (X-axis)(IUPred, (44)). **B)** N-protein undergoes concentration dependent LLPS with full-length gRNA. **C)** SARS-CoV-2 genome with regions tested for phase separation color coded: 5’-End (1-1000; turquoise), Frameshifting-region (13401-14400; magenta), homologous to published SARS-PS SARS-CoV packaging signal equivalent (19782-20363; green), Nucleocapsid fused to first 75 nt of 5’-End (grey line) (=subgenomic Nucleocapsid RNA; 1-75 + 28273-29533; purple) **D)** FRESCo (38) analysis of synonymous substitution restraints in ORF1ab. Significant synonymous constraints at four confidence cutoffs (1e-3, 1e-4, 1e-5, 1e-6) assessed over a ten-codon sliding window are marked by magenta lines. Tested regions correspond to those shown in C. **E)** Different RNA regions from SARS-CoV-2 (at 5nM) either drive or solubilize N-protein (1 µM) droplets. **F)** Ability of 5’-End and Frameshifting-region RNA to drive or solubilize condensation of N-protein (4 µM) over increasing RNA concentrations. **G)** Phase diagram of N-protein with either 5’-End or Frameshifting-region RNA at indicated concentrations. Quantification corresponds to microscopy images in Figure S2A. **H)** Length dependence of N-protein (2 µM) LLPS was assessed with Frameshifting-region and 5’-End RNAs extended with non-specific plasmid sequences (at 5 nM RNA). Scale bar, 8 µm unless otherwise noted.

### N-protein phase separates with viral RNA in a length, sequence and concentration dependent manner

We reconstituted purified N-protein under physiological buffer conditions with viral RNA segments and observed that N-protein produced in mammalian cells (post-translationally modified) or bacteria (unmodified) phase separated with viral RNA segments. However, unmodified protein yielded larger and more abundant droplets **(Figure S1A)**. Since N-protein in SARS-CoV1 virions is hypophoshorylated (*5*) and packaging (initiated by binding of N-protein to gRNA) first occurs in the cytoplasm of coronaviruses (*6, 7*), where N-protein is thought to be in its unphosphorylated state (*8*), we used unmodified protein for subsequent experiments.

Pure N-protein demixed into droplets on its own and phase separation was enhanced by full-length genomic SARS-CoV-2 RNA **(Figure 1B)**. To determine if certain segments of SARS-CoV-2 genome had preferential ability to drive phase separation, we identified regions of the gRNA under synonymous codon constraints. We hypothesized that LLPS occurs specifically with gRNA carrying a viral packaging signal(s), whose exact structure and location in corona viruses vary (*1, 9-12*) and is unknown for SARS-CoV-2. We identified multiple regions with reduced synonymous sequence substitutions, indicative of functional RNA sequences and structures **(Figure 1C and D, Figure S1B**). We focused on regions that also contained structures predicted to be conserved using RNAz (Table S1), located in ORF1ab RNA (which contains the packaging signals for other Betacorona viruses), occurs in packaged full-length genome sequences, and is absent from sub-genomic fragments (*13*) **(Figure 1C,D)**. We synthesized sequences corresponding to four regions: a region spanning the 5’-End (first 1000nts), the Frameshifting-region (1000 nts around the Frameshifting-element), a sequence corresponding to the SARS-CoV packaging signal, and the highly expressed subgenomic RNA sequence coding for the N-protein (containing the first 75 nucleotides of the 5’-UTR recombined onto the N-protein coding sequence) (*13*) **(Figure 1C)**.

N-protein LLPS varied as a function of the specific RNA-co-component **(Figure 1E)**. The 5’-End and the subgenomic Nucleocapsid-encoding RNAs promoted LLPS. In contrast, the Frameshifting-region and the SARS-CoV packaging signal-homologous region RNAs reduced phase separation relative to N-protein alone **(Figure 1E)**. The 5’-End and Frameshifting-region, which have the same length, displayed consistent near-opposing behaviors across a range of RNA and protein concentrations. The 5’-End generally drives N-protein condensation, whereas the Frameshifting-region solubilized condensates and promoted LLPS only within a narrow protein and RNA concentration range **(Figure 1F/G, Figure S1C)**. The concentration range at which the 5’-End could drive LLPS of N-protein was similar to the LLPS behavior of the full-length RNA genome **(Figures 1B, 1F)** in terms of total nucleic acid amount but not molar ratio.

N-protein binds gRNA in the cytosol in the presence of non-viral RNAs. We therefore assessed how non-viral, lung RNA influences LLPS. Total lung RNA did not alter N-protein only LLPS; in contrast, when combined with viral 5’-End RNA, total lung RNA enhanced condensate size, number, and viral RNA recruitment **(Figure S1D/E/F)**. gRNA is longer than many host RNAs and all subgenomic RNAs and we reasoned that length contributes to an electrostatically driven component of N-protein LLPS given protein’s pI is 10.07. Addition of 0.3 kb or 2.4 kb of non-viral sequence to the 1 kb 5’-End or Frameshift-region RNAs resulted in progressive LLPS enhancement, with the 5’-End driving greater LLPS relative to the Frameshifting-region at all lengths tested **(Figure 1H)**. In sum, N-protein undergoes LLPS under physiological conditions, including in the presence of abundant non-specific RNA, and LLPS is enhanced by viral RNA. Both specific viral RNA sequences and RNA length contribute to LLPS.

### Role of temperature and material properties in dictating N-protein condensation with viral RNA

Corona virus replication is most efficient at 33°C (*14*) and we therefore assessed the temperature-dependence of phase separation. N-protein alone demixed into droplets in a temperature-dependent manner, most pronounced at fever temperature (40°C) **(Figure 2A/B/C/D)**. Addition of the 5’-End RNA resulted in larger N-protein-containing droplets at 37 and 33°C (corresponding to lung and upper airway temperatures, respectively) (*15*). Similar results were obtained for N-protein/Nucleocapsid RNA condensates **(Figure S2A)**. This temperature behavior is notable both for its overlap with clinical features and because there are few biological polymers whose phase separation is enhanced at high temperature.

**Figure 2:**
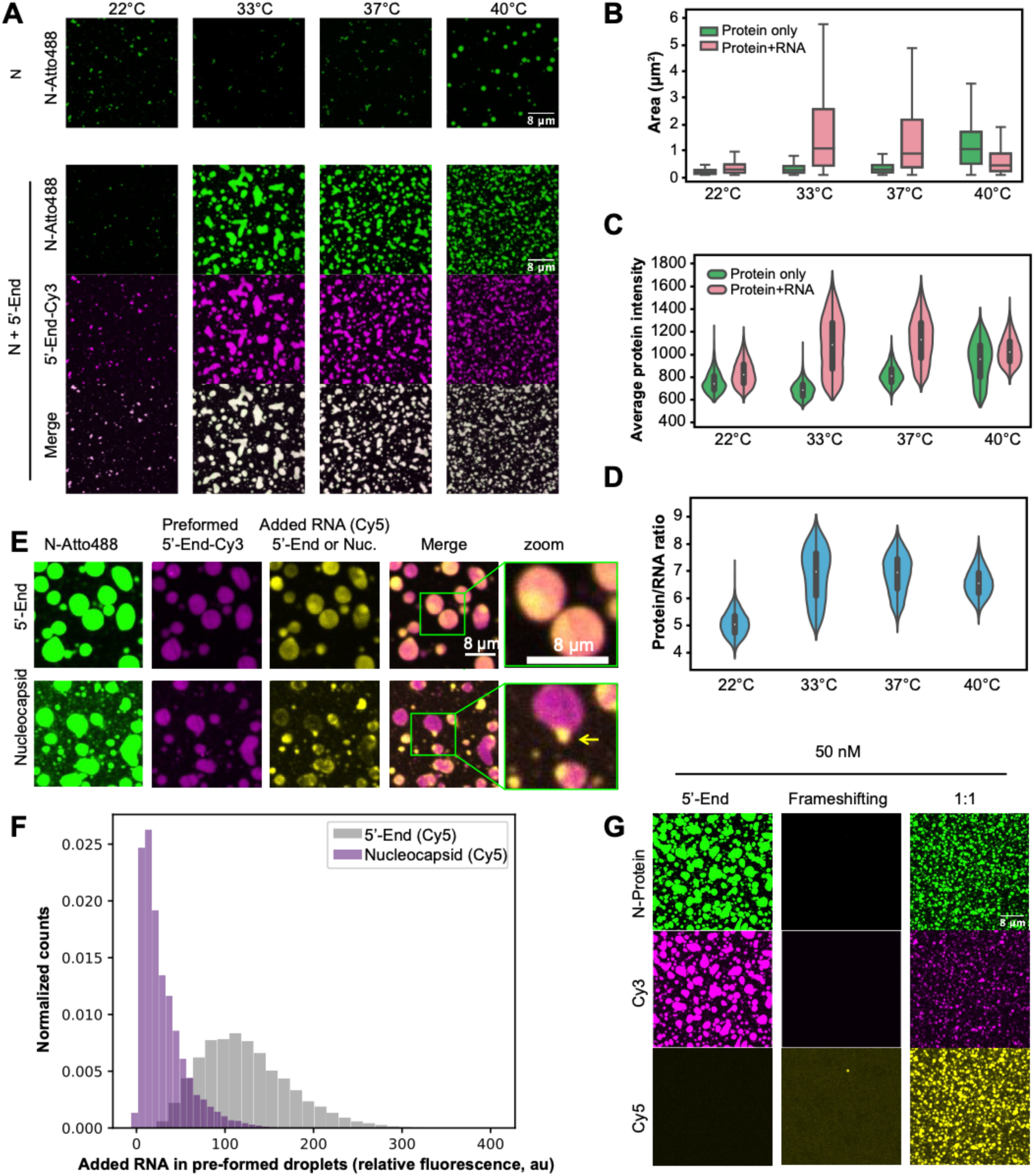
Temperature dependence and specificity of phase separation. **A)** N-protein (green) phase separates in a temperature-dependent manner (upper panel). Temperature-dependence is shifted when viral 5’-End RNA (magenta) is present (lower panel). **B)** Quantification of average protein intensity from (B) based on fluorescence intensity. **C)** Quantification of droplet area from (B). **D)** Quantification of protein/RNA ratio based on fluorescence intensity. **E)** 5’-End (yellow, upper panel) is recruited into preformed 5’-End /N-protein droplets (pink and green) but Nucleocapsid RNA (yellow, lower panel) is not efficiently recruited. **F)** Quantification of (E) showing intensity of second RNA added to preformed droplets. **G)** Mixing 5’-End and Frameshifting-region RNAs makes N-protein condensates with intermediate properties. Scale bar, 8 µm unless otherwise noted. Violin plots are scaled to have equal widths. Outliers not shown.

In infected cells, subgenomic viral RNAs, like Nucleocapsid RNA, are highly abundant RNA species (*13*). We hypothesized that material property differences contribute to selective packaging of gRNA and examined N-protein condensates made with RNAs that yielded different material properties. The 5’-End promoted larger, more liquid-like condensates; in contrast, the Nucleocapsid RNA and a non-viral (luciferase) RNA induced smaller, solid-like, flocculated condensates **(Figure S2B)**. To assess relevance of these material differences to selectivity, we added subgenomic Nucleocapsid RNA to preformed 5’-End droplets. Subgenomic Nucleocapsid RNA was excluded from preformed N-protein–5’-End condensates, whereas additional 5’-End RNA readily mixed **(Figure 2E/F)**. Thus, material properties of N-protein condensates have RNA sequence specificity that could act to exclude subgenomic RNAs from virions.

Different viral RNAs thus can promote or limit phase separation and yield different material properties in N-protein condensates **(Figure S2B)**. We hypothesized some RNA segments might function to maintain liquidity, and oppose problematic solidification in the context of long, full-length gRNA. Given the Frameshifting-region promoted dissolution, we examined whether this RNA could solubilize droplets made of other RNAs. We mixed Frameshifting-region with either 5’-End or Nucleocapsid RNA. Mixtures containing the 5’-End and Frameshifting-region produced droplets of intermediate properties, including smaller size. Similarly, Frameshifting-region RNA made Nucleocapsid RNA condensates less flocculated and more liquid-like **(Figure 2G, Figure S2C/D)**. These data suggest that distinct genomic RNA regions can promote or oppose phase separation and in combination may yield optimum material properties for packaging the whole genome.

### RNA sequence and structure attributes encode material properties

We next examined how viral RNA sequence and structure encode distinct LLPS behavior and droplet material properties. We experimentally assessed and modeled 5’-End and Frameshifting-region structures using SHAPE-MaP **(Figure 3A-D, Figure S3A/B)** (*16*). Both RNAs are highly structured. These RNA regions have similar fractions of SHAPE-reactive and -unreactive nucleotides and nucleotides modeled as base paired **(Figure S4D)**. However, the Frameshifting-Region forms a greater number of more complex multi-helix junction structures and has a higher A/U content (62% vs 52% for 5’-End) **(Figure 3A/B, Figure S4B/C)**. We next measured N-protein interactions with viral RNAs using RNP-MaP which selectively crosslinks lysine residues to proximal RNA nucleotides, largely independent of nucleotide identity and local RNA structure (**Figure S4D**) (*17*). We mapped N-protein interactions at protein:RNA ratios that promote either diffuse or condensed droplets for both the 5’-End and Frameshifting-region **(Figure S4A)**. For the 5’-End in the diffuse state (20x excess protein), there are two strong N-protein binding sites, and each occurs in a long A/U-rich unstructured region flanked by strong stem-loop structures **(Figure 3A/C**). In the droplet state (80x/160x excess protein) the two principal sites from the low-ratio state remain fully occupied and additional N-protein interaction sites appeared (the valency increased). In contrast, the Frameshifting-region showed generalized binding across the RNA by N-protein at all ratios **(Figure 3B/D)**. Binding was observed in both single-stranded regions and also in A/U rich structured regions **(Figure 3B/D**). In sum, N-protein interacts strongly with a few preferred sites in the 5’-End in both diffuse and condensed states, and interacts more homogeneously across the Frameshifting-region **(Figure 3F, Figure S4E)**.

**Figure 3:**
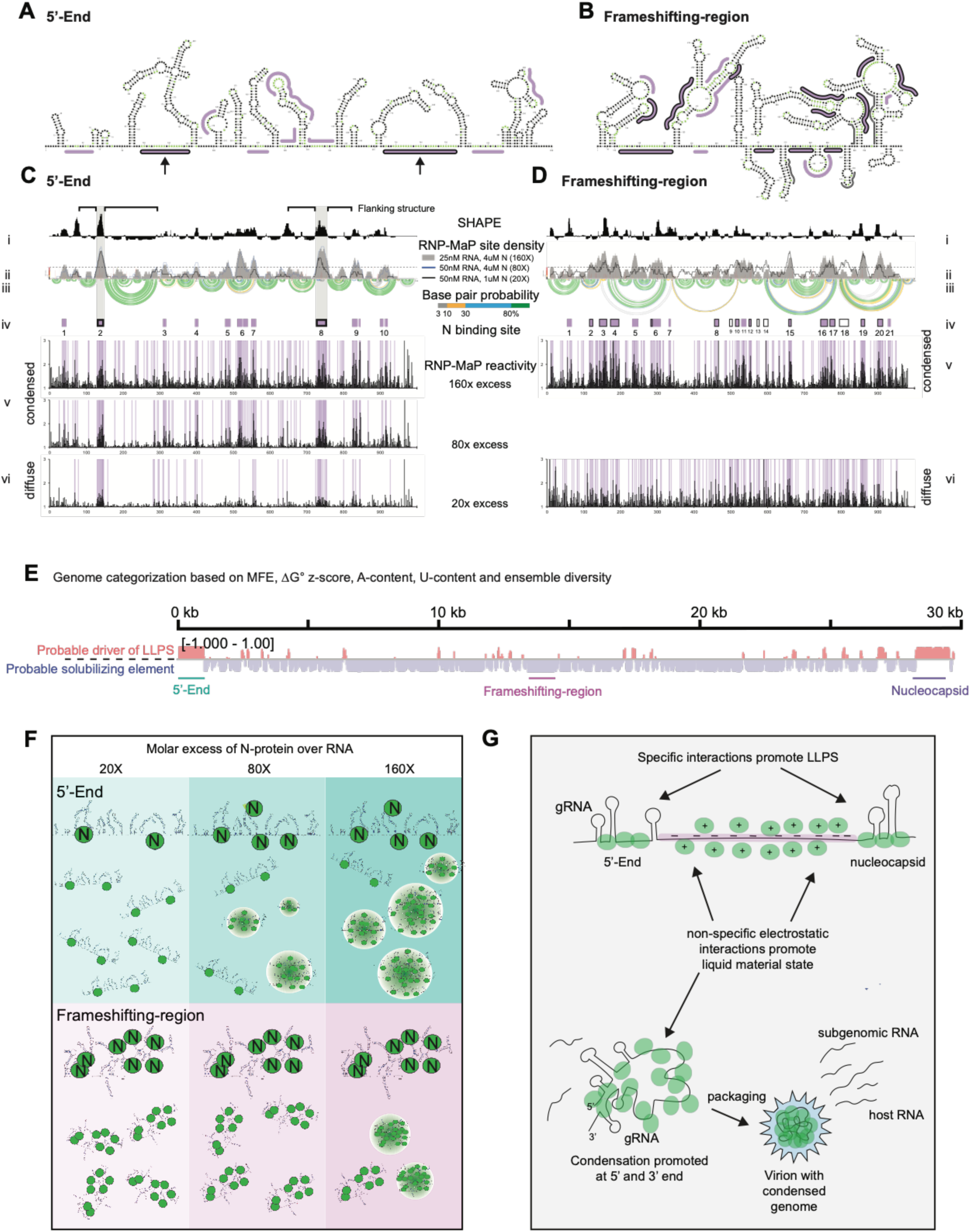
RNA sequence and structure encode interactions with N-protein. **A and B)** SHAPE-Map secondary structure models for the 5’-End (A) or Frameshifting-region (B). RNP-MaP N-protein binding sites, are purple. Arrows are two primary binding sites on the 5’-end RNA, both flanked by strong RNA structures emphasized with arrows. **C and D)** 5’-End (C) and Frameshift-region (D) display condition specific RNP-MAP reactivity in condensed and diffuse conditions. X-axis is the position in nucleotides, Y-axis is the reactivity (SHAPE or RNP-Map). i: SHAPE reactivity (black). ii: windowed average of RNP-MaP site density. Arcs indicate base pair probabilities (from SHAPE). iii: base pair probability. iv: N:protein binding site. (boxes: purple, at 160x; with black border, 20x). v-vi: Raw RNP-MaP reactivity in all conditions (15nt windows). Grey shading indicates positions of principle N: protein binding sites for 5’-End. Nucleotide RNP-MaP reactivities (black) are plotted for each RNA under 20x, 80x and 160x conditions with RNP-MaP sites (17). **E)** Genome similarity to 5’-End and Frameshifting-region. Mean for each feature is computed over all 120 base pair windows with center in the region of interest. MFE, dG z-score, and ensemble diversity are defined in (18). **F)** Model of for LLPS: Upper panel: 5’-End LLPS coincides with an increase in valency. Lower panel: frameshifting RNA has multiple binding sites that prevent condensate formation, unless N-protein excess is present to drive LLPS via protein-protein interaction. **G)** Model: packaging of gRNA may be a LLPS process driven by single-stranded regions flanked by structured regions (5’-End-like) that are stable N-protein binding sites. The majority of the genome resembles the solubilizing (Frameshifting-region-like), while the region coding for N-protein is similar to the 5’-End. The balance between LLPS-promoting and solubilizing elements may facilitate gRNA packaging.

We predicted the genome may be a mixture of sequences that promote LLPS like the 5’ End and that fluidize like the Frameshifting-region. Therefore, we computed structural properties of the 5’-End and Frameshifting-regions and compared to the rest of the gRNA. Most of the RNA genome has local minimum free energies for predicted structures, and A/U-content, ΔG z-score and ensemble diversity (ED) similar to that of the Frameshifting-region **(Figure 3E, Figure S4F/G)(*18*)**. Interestingly, the two major LLPS-promoting sequences, the 5’-End and Nucleocapsid-encoding region at the 3’ end of the genome **(Figure 1E)** share similar features **(Figure 3E/F/G, Figure S4F/G)**. In contrast, the internal gRNA is more similar to the Frameshifting-region and may act as a solubilizing element **(Figure 3F/G)** consistent with different regions of the genome having distinct contributions to LLPS of N-protein.

### N-protein phase separates in mammalian cells and can be disrupted by small molecules

To assess the ability of N-protein to phase separate in cells, we co-transfected HEK293 cells with N-protein fused to GFPspark and with H2B:mCherry (to mark nuclei in single cells). Cells with higher levels of transfection were more likely to form spherical droplets in the cytoplasm **(Figure 4A/B)**, suggesting N-protein condensation is concentration dependent. N-protein signal was generally excluded from the nucleus **(Figure 4C)**. N-protein droplets readily underwent fusion **(Figure 4D)** and recovered quickly following FRAP **(Figure 4E/F)** indicating dynamic recruitment of N-protein. These results suggest that N-protein can form cytoplasmic, liquid-like condensates in cells. Cells expressing N-protein may be used to screen for compounds that modify N-protein LLPS and thereby viral packaging.

**Figure 4:**
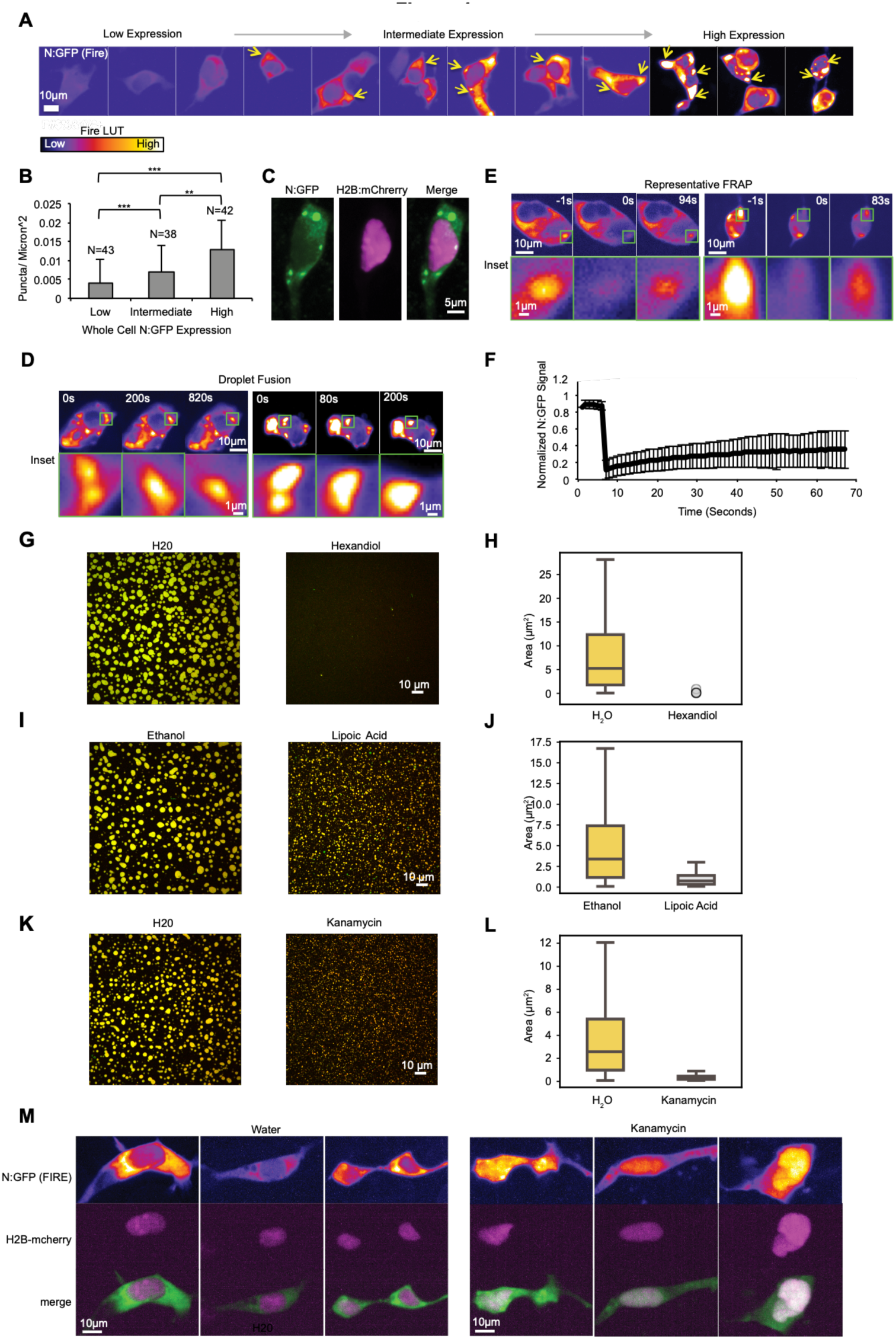
N-protein phase separates in mammalian cells and can be disrupted by small molecules. **A)** N-protein: GFP forms concentration dependent condensates in HEK293 cells. The fire LUT marks low expression in purple and high expression in yellow. **B)** Condensates per µ2 increased with N:GFP expression level. **C)** N:GFP was excluded from nuclei (marked with H2B:mCherry) of HEK293 cells. **D)** N:GFP condensates fused in HEK293 cells. Top panel: representative cells, Bottom panel: enlarged of fusion event, Scale bar, 1 µm. **E)** N:GFP condensates recovered partially after FRAP. Top panel shows representative condensate FRAP. Bottom panel: zoom of N:GFP condensate. Scale bar, 1 µm. **F)** Condensates recovered to 24% within 1 minute error bars show standard deviation from N=18 condensates. **G/H)** 9% 1-,6-hexanediole prevents N-protein/Frameshifting-region RNA LLPS. **I/J)** 1 mg/ml lipoic acid partially prevents N-protein/Frameshifting-region RNA LLPS. **K/L)** 5 mg/ml kanamycin partially prevents N-protein/Frameshifting-region RNA LLPS. M) 5 mg/ml kanamycin causes relocalization of N:GFP to the cell nucleus in 37% of treated cells (N= 105, 0% in H2O, N=100). Scale bar, 10 µm unless otherwise noted.

We reasoned that small molecules that increased or decreased N-protein LLPS or altered RNA recruitment could slow viral infection. We examined 1,6-hexanediol (*19*), lipoic acid (*20*), the aminoglycoside kanamycin (*21*) and cyclosporin (*22*), each of which potentially alters LLPS by a representative and distinctive mechanism. As a simple positive test, we examined 1,6-Hexanediol which disrupts LLPS (*19*). Indeed, 1,6-Hexanediol prevented condensate formation **(Figure 4G/H)**. Lipoic acid dissolves cellular stress granules (*20*), which are impacted in SARS-CoV (*23*). Lipoic acid treatment resulted in smaller condensates than vehicle **(Figure 4I/J)**. The aminoglycoside kanamycin binds promiscuously to nucleic acids via electrostatic interactions and, is an interesting candidate to disrupt LLPS due to its conformational adaptability and thus low RNA target specificity and was implicated as antiviral in HIV-1 by preventing RNA-protein interactions (*21*). Addition of non-toxic concentrations of kanamycin to droplets decreased the size of condensates in the reconstituted assay **(Figure 4K/L)** and N-protein relocalized to the nucleus in 37% of treated cells **(Figure 4M**).

An alternative to targeting N-protein–RNA LLPS directly is to target interactions between LLPS components and host proteins. We used a machine learning approach (*24*) to identify binding sites for host proteins on the SARS-Cov-2 RNA genome and identified PPIL4, an RNA-binding protein containing a prolyl isomerase motif **(Figure S5A)** and a putative target of cyclosporin. Intriguingly, cyclosporin was also suggested as an antiviral for coronaviruses (*22*) and another cyclophilin, CYPA, binds the nucleocapsid of SARS-CoV. Cells treated with cyclosporin more readily formed condensates (**Figure S5B/C)**. Prolyl isomerase activity within N-protein-mediated condensates may facilitate protein folding given the presence of 26 prolines in N-protein or may alter the nature of N-protein–RNA interactions.

## Discussion

A role of LLPS in SARS-CoV-2 is not only shown here, but is being independently confirmed and thus supported by multiple other research teams (Young, Yildiz, Soranno, Holehouse, Hyman groups private communication and Fawzi group simultaneous publication: https://www.biorxiv.org/content/10.1101/2020.06.09.141101v1.full.pdf). SARS-CoV-2 N-protein phase separates in an RNA sequence, length, and likely structure dependent manner (Figure 3F/G: model) and distinct regions of the viral RNA genome can either promote LLPS (5’-End region) or act as solubilizing elements (Frameshifting-region). Multivalent polymer interactions are a driving force of LLPS and we propose that the well-defined single-stranded sequences preferentially bound by N-protein enable the 5’-End region and the 3’-End regions, to promote specific LLPS. The Frameshifting-region, conversely, is more uniformly bound by N-protein in both diffuse and droplet states. Thus, the Frameshifting-region, and potentially the majority of the genome, may be coated by N-protein contributing non-specific electrostatic interactions that ensure fluidity of the large gRNA molecule. In this model, the full-length gRNA consists of a mixture of phase separation-promoting and aggregation-dissolving elements to promote regulated, selective LLPS. In line with our findings, recent work from the Holehouse and Soranno labs proposes a model in which a small number of high-affinity sites drives symmetry breaking to drive the assembly of single-RNA condensates instead of large multi-RNA droplet, providing a putative mechanism through which individual genomes could be packaged.

The temperature dependence for N-protein phase-behavior overlaps body temperatures from 33°C (temperature of upper airways) to 37°C and 40°C (fever), with increasing temperatures also increasing the propensity of RNA and protein to phase separate. Upper airway temperatures induce large and protein-rich condensates, suggestive of differential consequences on nucleocapsid LLPS and thereby SARS-CoV-2 biology in different parts of the body and at different stages of disease progression, including fever (*25*). SARS-CoV-2 infectivity in nasal epithelium shows a gradient in infectivity that correlates with the temperatures of maximum N-protein LLPS (*26*). The temperature-dependent LLPS of N-protein may also be relevant to virus incubation in reservoir species such as Chinese horseshoe bats (*27*). Cool body temperatures of bat hibernation may limit N-protein LLPS, slowing viral replication, while body warming following hibernation may induce LLPS and promote viral packaging and infection. As of yet, there are very few biological examples of phase separation characterized by lower critical solution temperature (LCST) (*28-30*), making LCST phase separation by N-protein an important system for understanding biological biomolecular condensation.

LLPS is an attractive target for compound screening because viral replication may be inhibited by molecules that either block or modify phase separation. The three compounds we tested for effects in vitro modulate LLPS in different ways, including impacting weak hydrophobic and protein-protein interactions (lipoic acid and hexanediol) or protein-RNA interactions (kanamycin). Specific RNA sequences and structures which regulate N-protein LLPS may also be targeted directly in the development of antiviral therapies. These straightforward in vitro and in vivo assays comprise a powerful starting point for evaluating FDA-approved compounds to reveal new classes of antiviral compounds that target phase-separation.

## Materials and Methods

### In vitro transcription

was carried out according to our established protocols (*31*). Orf1ab templates were synthesized (IDT) and cloned into pJet (ThermoFisher Scientific K1231) using blunt end cloning. Directionality and sequence were confirmed using Sanger sequencing (GENEWIZ). Plasmid were linearized with XBAI restriction enzyme (NEB R0145S) and gel purified (QIAGEN 28706). Nucleocapsid RNA was produced from pu57 Nucleocapsid, a kind gift from the Sheahan lab, linearized with NOTI (NEB R3189S) and STUI (NEB R0187S). 100 ng of gel purified DNA was used as a template for in vitro transcription (NEB E2040S) carried out according to the manufacturer’s instructions with the addition of 0.1µl of Cy3 (Sigma PA53026) or Cy5 (Sigma PA55026) labeled UTP to each reaction. Following incubation at 37°C for 18 hours, in vitro transcription reactions were treated with DNAseI (NEB M0303L) according to the manufacturer’s instructions. Following DNAse treatment, reactions were purified with 2.5M LiCL precipitation. Purified RNA amounts were quantified using nanodrop and verified for purity and size using a denaturing agarose gel and Millenium RNA ladder (ThermoFisher Scientific AM7151).

### Purification of genomic SARS-CoV-2 RNA (gRNA)

Vero E6 cells were cultured to ∼90% confluence in T175 flasks. Immediately prior to infection the culture medium was aspirated and cells were washed with PBS. Flasks were infected at a multiplicity of infection of 3 with SARS-CoV-2 at 37°C for 1 h. After 1 hour cells were supplemented with pre-warmed DMEM (Gibco) with 5% FetalCloneII (HyClone) and 1x Anti-Anti (Gibco). Cells were then incubated for an additional 24 hours at 37°C. After the infection was complete, the cell supernatant was aspirated and concentrated using Millipore Centrifugal Amicon filters to approximately 4 mL total volume. The supernatant was then lysed in TRIzol LS, and viral RNA was extracted from the trizol using chloroform extraction.

### Recombinant Protein Expression and Purification

For protein purification, full-length N-protein was tagged with an N-terminal 6-Histidine tag (pET30b-6xHis-TEV-Nucleocapsid,) and expressed in BL21 E. coli (New England Biolabs). All steps of the purification after growth of bacteria were performed at 4°C. Cells were lysed in lysis buffer (1.5M NaCl, 20 mM Phosphate buffer pH 7.5, 20 mM Imidazole, 10mg/mL lysozyme, 1 tablet of Roche EDTA-free protease inhibitor cocktail Millipore Sigma 11873580001) and via sonication. The lysate was then clarified via centrifugation (SS34 rotor, 20,000 rpm 30 minutes) and the supernatant was incubated and passed over a HisPur™ Cobalt Resin (ThermoFisher Scientific 89965) in gravity columns. The resin was then washed with 4X 10 CV wash buffer (1.5M NaCl, 20 mM Phosphate buffer pH 7.5, 20 mM Imidazole) and protein was eluted with 4 CV Elution buffer (0.25 M NaCl, 20 mM Phosphate buffer pH 7.5, 200 mM Imidazole). The eluate was then dialyzed into fresh storage buffer (0.25 M NaCl, 20 mM Phosphate buffer) and aliquots of protein were flash frozen and stored at -80 °C. Protein was checked for purity by running an SDS-PAGE gel followed by Coomassie staining as well as checking the level of RNA contamination via Nanodrop and through running of a native agarose RNA gel. Please note, that while self-purified protein had very low RNA contamination, commercially acquired bacterial expressed N-protein at similar concentrations had a high contamination of RNA that severely enhanced LLPS. This enhancement of LLPS was abrogated through addition of RNaseA (Qiagen 19101).

### Dyeing of N-protein

N-protein was dyed by adding (3:1) Atto 488 NHS ester (Millipore Sigma 41698) to purified protein and incubating mix at 30°C for 30 minutes. Unbound dye was removed by 2 washes with 100X excess of protein storage buffer followed by centrifugation in Amicon® Ultra-4 Centrifugal Filter Units (SIGMA MilliPORE). LLPS of dyed and undyed protein was compared as quality control and results were similar (data not shown).

### Phase separation assays

For in vitro reconstitution LLPS experiments, 15 µl droplet buffer (20 mM Tris pH 7.5, 150 nM NaCl) was mixed with cy3 or cy5 labeled desired RNA and 5 µl protein in storage buffer was added at desired concentration. The mix was incubated in 384-well plates (Cellvis P384-1.5H-N) for 16 hours at 37°C unless indicated otherwise. Droplets already formed after short incubations of 20 minutes or less, however, they were initially smaller and matured into larger droplets during the overnight incubation step. Imaging of droplets was done on a spinning disc confocal microscope (Nikon CSU-W1) with VC Plan Apo 100X/? NA oil (Cargille Lab 16241) immersion objective and an sCMOS 85% QE camera (Photometrics). Data shown are representative of three or more independent replicates, across several RNA preparations.

### Sequestration experiments

For sequestration experiments, N-protein/5’-End (Cy3) condensates were preformed and after 1.5 h incubation, 5 nM cy5-labeled RNA of interest was added mixed, and incubated for another 14 hours before imaging.

### Drug treatments of in vitro phase separation assays

For drug treatment of in vitro phase separation assays, droplet buffer was pre-mixed with drugs or vehicles, before RNA and protein were added to the mix. (R)-(+)-α-Lipoic acid (Sigma-Aldrich, cat. number: 07039) was added at 1 mg/ml in the presence of excess DTT to reduce its thiole ring and compared to the vehicle ethanol. 1,6-Hexanediol (Sigma Aldrich, cat. number: 240117) was added to a final concentration of 9%. Kanamycin (Millipore Sigma 60615-25G) was added to a final concentration of 5 mg/ml. 1,6-Hexanediol and Kanamycin were compared to the vehicle H20. We chose this order of component addition, to most closely mimic possible screening conditions, in which drugs would likely be pre-added to multi-well plates. The mixtures were then incubated for 16 h at 37°C before imaging.

### RNP-MaP probing of N-Protein-RNA interactions

N-Protein and RNA mixtures were prepared as described in the “Phase Separation Assay” section above and incubated for 1.5 hours at 37°C. N-Protein–5′-End RNA mixtures were prepared in three conditions: (1) 50nM RNA, 1µM protein (diffuse state, 20x excess protein), (2) 50nM RNA, 4µM protein (droplet state, 80x excess protein), and (3) 25nM RNA, 4µM protein (droplet state, 160x excess protein). N-Protein–Frameshifting-region RNA mixtures were prepared in two conditions: (1) 50nM RNA, 1µM protein (diffuse state, 20x excess protein) and (2) 25nM RNA, 4µM protein (droplet state, 160x excess protein). RNA-only samples were also prepared as a control. After confirmation of phase separation by imaging **(Figure S4A)** mixtures were immediately subjected to RNP-MaP treatment as described (*17*), with modifications described below. Briefly, 200 µl of mixtures were added to 10.5 µl of 200 mM SDA (in DMSO) in wells of a 6-well plate and incubated in the dark for 10 minutes at 37°C. RNPs were crosslinked with 3 J/cm^2^ of 365 nm wavelength UV light. To digest unbound and crosslinked N-proteins, reactions were adjusted to 1.5% SDS, 20 mM EDTA, 200mM NaCl, and 40mM Tris-HCl (pH 8.0) and incubated at 37°C for 10 minutes, heated to 95°C for 5 minutes, cooled on ice for 2 minutes, and warmed to 37°C for 2 minutes. Proteinase K was then added to 0.5 mg/ml and incubated for 1 hour at 37°C, followed by 1 hour at 55°C. RNA was purified with 1.8× Mag-Bind TotalPure NGS SPRI beads (Omega Bio-tek), purified again (RNeasy MinElute columns, Qiagen), and eluted with 14 µl of nuclease-free water.

### SHAPE-MaP RNA structure probing

SHAPE-MaP treatment with 5NIA was performed as described (*32*). Briefly, *In* vitro transcribed 5’-End or Frameshifting-region RNA (1200 ng in 40µL nuclease-free water) was denatured at 95°C for 2 minutes followed by snap cooling on ice for 2 minutes. RNA was folded by adding 20 µL of 3.3× SHAPE folding buffer [333mM HEPES (pH 8.0), 333mM NaCl, 33mM MgCl2] and incubating at 37°C for 20 minutes. RNA was added to 0.1 volume of 250 mM 5NIA reagent in DMSO (25 mM final concentration after dilution) and incubated at 37°C for 10 minutes. No-reagent (in neat DMSO) control experiments were performed in parallel. After modification, all RNA samples were purified using RNeasy MiniElute columns and eluted with 14 µl of nuclease-free water.

### MaP reverse transcription

After SHAPE and RNP-MaP RNA modification and purification, MaP cDNA synthesis was performed using a revised protocol as described (*33*). Briefly, 7 µL of purified modified RNA was mixed with 200 ng of random 9-mer primers and 20 nmol of dNTPs and incubated at 65°C for 10 min followed by 4°C for 2 min. 9 µL 2.22× MaP buffer [1× MaP buffer consists of 6 mM MnCl2, 1 M betaine, 50 mM Tris (pH 8.0), 75 mM KCl, 10 mM DTT] was added and the combined solution was incubated at 23°C for 2 min. 1 μL SuperScript II Reverse Transcriptase (200 units, Invitrogen) was added and the reverse transcription (RT) reaction was performed according to the following temperature program: 25°C for 10 min, 42°C for 90 min, 10×[50°C for 2 min, 42°C for 2 min], 72°C for 10 min. RT cDNA products were then purified (Illustra G-50 microspin columns, GE Healthcare).

### Library preparation and Sequencing

Double-stranded DNA (dsDNA) libraries for sequencing were prepared using the randomer Nextera workflow (*34*). Briefly, purified cDNA was added to an NEBNext second-strand synthesis reaction (NEB) at 16°C for 150 minutes. dsDNA products were purified and size-selected with SPRI beads at a 0.8× ratio. Nextera XT (Illumina) was used to construct libraries according to the manufacturer’s protocol, followed by purification and size-selection with SPRI beads at a 0.65× ratio. Library size distributions and purities were verified (2100 Bioanalyzer, Agilent) and sequenced using 2×300 paired-end sequencing on an Illumina MiSeq instrument (v3 chemistry).

### Sequence alignment and mutation parsing

FASTQ files from sequencing runs were directly input into *ShapeMapper 2* software (*35*) for read alignment, mutation counting, and SHAPE reactivity profile generation. The *--random-primer-len 9* option was used to mask RT primer sites with all other values set to defaults. For RNP-MaP library analysis, the protein:RNA mixture samples are passed as the *--modified* samples and no-protein control RNA samples as *-- unmodified* samples. Median read depths of all SHAPE-MaP and RNP-MaP samples and controls were greater than 50,000 and nucleotides with a read depth of less than 5000 were excluded from analysis.

### Secondary structure modeling

The *Superfold* analysis software (*34*) was used with SHAPE reactivity data to inform RNA structure modeling by *RNAStructure* (*36*). Default parameters were used to generate base-pairing probabilities for all nucleotides (with a max pairing distance of 200 nt) and minimum free energy structure models. The local median SHAPE reactivity were calculated over centered sliding 15-nt windows to identify structured RNA regions with median SHAPE reactivities below the global median. Secondary structure projection images were generated using the (VARNA) visualization applet for RNA (*37*).

### RNP-MaP reactivity analysis

A custom RNP-MaP analysis script (*17*) was used to calculate RNP-MaP “reactivity” profiles from the *Shapemapper 2* “profile.txt” output. RNP-MaP “reactivity” is defined as the relative MaP mutation rate increase of the crosslinked protein-RNA sample as compared to the uncrosslinked (no protein control) sample. Nucleotides whose reactivities exceed reactivity thresholds are defined as “RNP-MaP sites”. RNP-MaP site densities were calculated over centered sliding 15-nt windows to identify RNA regions bound by N-protein. An RNP-MaP site density threshold of 5 sites per 15-nt window was used to identify “N-protein binding sites” with boundaries defined by the RNP-MaP site nucleotides.

## Mammalian cells methods

### Cell Culture

HEK293 cells were originally obtained from ATCC. HEK293 cells were maintained in DMEM (Corning 10-013-CV) supplemented with 10% Fetal Bovine Serum (Seradigm V500-050). No antibiotics were used.

### Transfection

24 hours prior to transfection, confluent cells were split 1:5. Two hours prior to transfection, 2mL of fresh media was added to 10cm dishes. 25ug of plasmid DNA for each Nucleocapsid Spark (Sino biological VG40588-ACGLN) and H2BmCherry (from Jun Lu lab Yale University) was co-transfected using calcium phosphate. Following transfection, cells were incubated for 72 hours prior to imaging or drug treatment.

### Drug Treatment

Cells were incubated with drugs for 24 hours prior to imaging. Kanamycin (Millipore Sigma 60615-25G) was added to a final concentration of 5 mg/ml with control cells treated with 10% sterile water by volume or 0.001mg/ml cyclosporin A (Millipore Sigma SML1018-1ML) and control 0.001% DMSO vehicle.

## Microscopy and image analysis

### Cell Imaging

Cells were imaged using a 40X air objective on a spinning disk confocal microscope (Nikon Ti-Eclipse, Yokogawa CSU-X1 spinning disk). Images were taken with a Photometrics Prime 95B sCMOS camera. Representative cells are taken from at least 6 biological replicates pooled from at least 3 independent rounds of transfection/drug treatment.

### Analysis of cell imaging data

Figures A/E/D/M (top) depict maximum intensity projections with Fire LUTs. Average fluorescence intensity and area were obtained by thresholding max projections in ImageJ. Number of puncta per cell was manually counted from max projections.

### FRAP

Prior to bleaching, cells were imaged for seven seconds with one second per frame. Following bleaching with 488 nm laser, puncta recovery was monitored for at least one minute with one second frame intervals. Puncta fluorescence recovery was quantified using image J. Fluorescence was normalized by subtracting background fluorescence and relative to both the initial unbleached signal and an unbleached puncta in an unbleached cell in the same frame. Quantification represents 18 puncta from 15 movies with error bars depicting standard deviation.

### Quantification of in vitro condensates

For each image, droplets were segmented based on a threshold of 4*background intensity. Any segmented region with an area less than 0.07 μm^2^ was removed. The average protein and RNA intensity values within each droplet were calculated, and protein/RNA ratios were determined by dividing these averages on a per-droplet basis. For protein intensity and area, a two-sample Kolmogorov-Smirnov (KS) test was applied to compare protein-only with protein+RNA distributions at each temperature. A two-sample KS test was also used to make pairwise comparisons between each of the protein-only distributions, and between each of the protein+RNA distributions. Similarly, a two-sample KS test was performed to compare protein/RNA ratios. Images were processed in ImageTank [cite] and plotted with Python using Matplotlib and Seaborn. Statistics were performed in Python with SciPy.

### Quantification of colocalization

For co-localization analysis, cy5 intensity (signal for 5’-End or Nucleocapsid RNA) was plotted for pixels with >2X background cy3 intensity (signal for 5’-End RNA). Values were represented by a histogram of cy5 intensity.

## Computational sequence analysis

### FRESCo: Detection of regions of excess synonymous constraint

To detect regions of excess synonymous constraint, we used the FRESCo framework (*38*) which detects mutational differences between strains taking into account overlapping features. We scanned for genic translated regions for excess constraint at 10 1, 5, 10, 20, and 50 codon sliding windows. The regions of synonymous constraint were detected based on a set of 44 Sarbecovirus genomes listed in (*39*). Genic regions were extracted, translated, and aligned based on the amino acid sequence using Muscle version 3.8.31 (*40*). For each gene, sequences with less than 25% identity to the reference SARS-CoV-2 sequence (NC_045512) were removed. A nucleotide-level codon alignment was constructed based on the amino acid alignment, and gene-specific phylogenetic trees were constructed using RAxML version 8.2.12 with the GTRGAMMA model of nucleotide evolution (*41*). Regions with excess synonymous constraint at a significance level of 1e-5 in ten codon windows were extracted for further analysis. Thirty base pairs of flanking sequence were added on either side of each synonymous constraint element and RNAz 2.1 (*42*) was used to scan for conserved, stable RNA structures. The rnazWindow.pl script was used to filter alignments and divide into 120 base pair windows. Secondary structure detection was performed for both strands with SVM RNA-class probability set to 0.1.

### Seqweaver: Mapping of RBP-RNA interactions

To map RBP-RNA interactions across the SARS-CoV-2 viral RNA genomes, we leverage a deep learning-based sequence model, Seqweaver (*24*), whose accuracy in predicting RBP target site regulation we previously extensively evaluated. We split the SARS-CoV-2 viral genomes into 10 bp bins, and predicted the RBP binding probability for each bin while taking into account the 1,000bp flanking viral RNA sequence context. To identify robust viral binding sites, we transformed the Seqweaver predicted probability into Z-scores, where the background distribution was set to the human host transcriptome RBP binding site probability distribution mapped via CLIP-seq (*43*).

### Genome Analysis

A support vector machine with a linear kernel was trained using the scikit-learn Python library to distinguish between 120-base pair sliding windows in the 5’-End and the Frameshifting-region of SARS-CoV-2 (NC_045512.2) based on the following features: percent A content, percent U content, mean free energy ΔG, ΔG Z-score, and ensemble diversity, where mean free energy ΔG, ΔG Z-score, and ensemble diversity values were taken from (*18*). Features were scaled before classification to have mean of 0 and standard deviation of 1. All windows with center <= 1000 were used for the 5’-End, and all windows with center >= 13401 and <14401 were used for the Frameshifting-region. The classifier was then applied to all 120 bp windows outside the 5’-End and Frameshifting-region. The probability estimate for each sliding window of assignment to the 5’-End was plotted, after linearly re-scaling the probabilities for visualization purposes to have maximum of 1 and minimum of -1. Windows in the 5’-End are plotted with their class labels of 1, and windows in the FSE are plotted with their class labels of -1.

### Plasmids and sequences used

To create pET30b-6xHis-TEV-Nucleocapsid, the N-protein coding sequence (28273-29533 nt) preceded by the ORFN subgenomic 5’-UTR (1-75 nt) (Plasmid pUC57-2019-ncov, kind gift from Tim Sheahan and Ralph Baric) was cloned into AGB1329 (pET30b-6xHis-TEV) using SALI (NEB R3138S) and NOTI (NEB R3189S) restriction cloning with addition of the restriction sites to pUC57-2019-ncov by PCR using Sal-N-protein-fw (acgcatcgtcgacATGTCTGATAATGGACC-CCAAAATCAG) and Not1-N-protein-rev (tatctatgcggccgcTTAGGCCTGAGTTGAGTCAGC). SARS-CoV-2 genome regions for templates of RNA production (5’-End (1-1000 nt), Frameshifting-region (13401-14400 nt), SARS-CoV equivalent PS (19782-20363 nt) were derived from MN988668.1 Severe acute respiratory syndrome coronavirus 2 isolate 2019-nCoV WHU01. Sequence of N-protein for purification is as follows. atgcaccatcatcatcatcattcttctggtgaaaacctgtattttcagggcgtcgacATGTCTGATAATGGACCCCAAAATCAGCGAAATGCACCCCGCATTACGTTTGGTGGACCCTCAGATTCAACTGGCAGTAACCAGAATGGAGAACGCAGTGGGGCGCGATCAAAACAACGTCGGCCCCAAGGTTTACCCAATAATACTGCGTCTTGGTTCACCGCTCTCACTCAACATGGCAAGGAAGACCTTAAATTCCCTCGAGGACAAGGCGTTCCAATTAACACCAATAGCAGTCCAGATGACCAAATTGGCTACTACCGAAGAGCTACCAGACGAATTCGTGGTGGTGACGGTAAAATGAAAGATCTCAGTCCAAGATGGTATTTCTACTACCTAGGAACTGGGCCAGAAGCTGGACTTCCCTATGGTGCTAACAAAGACGGCATCATATGGGTTGCAACTGAGGGAGCCTTGAATACACCAAAAGATCACATTGGCACCCGCAATCCTGCTAACAATGCTGCAATCGTGCTACAACTTCCTCAAGGAACAACATTGCCAAAAGGCTTCTACGCAGAAGGGAGCAGAGGCGGCAGTCAAGCCTCTTCTCGTTCCTCATCACGTAGTCGCAACAGTTCAAGAAATTCAACTCCAGGCAGCAGTAGGGGAACTTCTCCTGCTAGAATGGCTGGCAATGGCGGTGATGCTGCTCTTGCTTTGCTGCTGCTTGACAGATTGAACCAGCTTGAGAGCAAAATGTCTGGTAAAGGCCAACAACAACAAGGCCAAACTGTCACTAAGAAATCTGCTGCTGAGGCTTCTAAGAAGCCTCGGCAAAAACGTACTGCCACTAAAGCATACAATGTAACACAAGCTTTCGGCAGACGTGGTCCAGAACAAACCCAAGGAAATTTTGGGGACCAGGAACTAATCAGACAAGGAACTGATTACAAACATTGGCCGCAAATTGCACAATTTGCCCCCAGCGCTTCAGCGTTCTTCGGAATGTCGCGCATTGGCATGGAAGTCACACCTTCGGGAACGTGGTTGACCTACACAGGTGCCATCAAATTGGATGACAAAGATCCAAATTTCAAAGATCAAGTCATTTTGCTGAATAAGCATATTGACGCATACAAAACATTCCCACCAACAGAGCCTAAAAAGGACAAAAAGAAGAAGGCTGATGAAACTCAAGCCTTACCGCAGAGACAGAAGAAACAGCAAACTGTGACTCTTCTTCCTGCTGCAGATTTGGATGATTTCTCCAAACAATTGCAACAATCCATGAGCAGTGCTGACTCAACTCAGGCCTAA

Complete RNA sequences are as follows.

T7 Plasmid sequences

Frameshifting-region

GGGAGAGCGGCCGCCAGATCTTCCGGATGGCTCGAGTTTTTCAGCAAGATTGGCTGTAGTTGTGATCAACTCCGCGAACCCATGCTTCAGTCAGCTGATGCACAATCGTTTTTAAACGGGTTTGCGGTGTAAGTGCAGCCCGTCTTACACCGTGCGGCACAGGCACTAGTACTGATGTCGT ATACAGGGCTTTTGACATCTACAATGATAAAGTAGCTGGTTTTGCTAAATTCCTAAAAACTAATTGTTGTCGCTTCCAAGAAAAGGACGAAGATGACAATTTAATTGATTCTTACTTTGTAGTTAAGAGACACACTTTCTCTAACTACCAACATGAAGAAACAATTTATAATTTACTTAAGGATTGTCCAGCTGTTGCTAAACATGACTTCTTTAAGTTTAGAATAGACGGTGACATGGTACCACATATATCACGTCAACGTCTTACTAAATACACAATGGCAGACCTCGTCTATGCTTTAAGGCATTTTGATGAAGGTAATTGTGACACATTAAAAGAAATACTTGTCACATACAATTGTTGTGATGATGATTATTTCAATAAAAAGGACTGGTATGATTTTGTAGAAAACCCAGATATATTACGCGTATACGCCAACTTAGGTGAACGTGTACGCCAAGCTTTGTTAAAAACAGTACAATTCTGTGATGCCATGCGAAATGCTGGTATTGTTGGTGTACTGACATTAGATAATCAAGATCTCAATGGTAACTGGTATGATTTCGGTGATTTCATACAAACCACGCCAGGTAGTGGAGTTCCTGTTGTAGATTCTTATTATTCATTGTTAATGCCTATATTAACCTTGACCAGGGCTTTAACTGCAGAGTCACATGTTGACACTGACTTAACAAAGCCTTACATTAAGTGGGATTTGTTAAAATATGACTTCACGGAAGAGAGGTTAAAACTCTTTGACCGTTATTTTAAATATTGGGATCAGACATACCACCCAAATTGTGTTAACTGTTTGGATGACAGATGCATTCTGCATTGTGCAAACTTTAATGTTTTATTCTCTACAGTGTATC TTT

5’-End

T7 Plasmid sequences

5’-UTR ORF1ab

5’-UTR recombined onto Nucleocapsid

GGGAGAGCGGCCGCCAGATCTTCCGGATGGCTCGAGTTTTTCAGCAAGATTTAAAGGTTTATACCTTCCCAGGTAACAAACCAACCAACTTTCGATCTCTTGTAGATCTGTTCTCTAAACGAACTTTAAAATCTGTGTGGCTGTCACTCGGCTGCATGCTTAGTGCACTCACGCAGTATAATTAATAACTAATTACTGTCGTTGACAGGACACGAGTAACTCGTCTATCTTCTGCAGGCTGCTTACGGTTTCGTCCGTGTTGCAGCCGATCATCAGCACATCTAGGTTTCGTCCGGGTGTGACCGAAAGGTAAGATGGAGAGCCTTGTCCCTGGTTTCAACGAGAAAACACACGTCCAACTCAGTTTGCCTGTTTTACAGGTTCGCGACGTGCTCGTACGTGGCTTTGGAGACTCCGTGGAGGAGGTCTTATCAGAGGCACGTCAACATCTTAAAGATGGCACTTGTGGCTTAGTAGAAGTTGAAAAAGGCGTTTTGCCTCAACTTGAACAGCCCTATGTGTTCATCAAACGTTCGGATGCTCGAACTGCACCTCATGGTCATGTTATGGTTGAGCTGGTAGCAGAACTCGAAGGCATTCAGTACGGTCGTAGTGGTGAGACACTTGGTGTCCTTGTCCCTCATGTGGGCGAAATACCAGTGGCTTACCG CAAGGTTCTTCTTCGTAAGAACGGTAATAAAGGAGCTGGTGGCCATAGTTACGGCGCCGATCTAAAGTCATTTGACTTAGGCGACGAGCTTGGCACTGATCCTTATGAAGATTTTCAAGAAAACTGGAACACTAAACATAGCAGTGGTGTTACCCGTGAACTCATGCGTGAGCTTAACGGAGGGGCATACACTCGCTATGTCGATAACAACTTCTGTGGCCCTGATGGCTACCCTCTTGAGTGCATTAAAGACCTTCTAGCACGTGCTGGTAAAGCTTCATGCACTTTGTCCGAACAACTGGACTTTATTGACACTAAGAGGGGTGTATACTGCTGCCGTGAACATGAGCATGAAATTGCTTGGTACACGGAACGTTCTATCTTT

SARS-CoV2 (homologous to SARS-CoV Packaging signal)

GGGAGAGCGGCCGCCAGATCTTCCGGATGGCTCGAGTTTTTCAGCAAGATTTTGAGCTTTGGGCTAAGCGCAACATTAAACCAGTACCAGAGGTGAAAATACTCAATAATTTGGGTGTGGACATTGCTGCTAATACTGTGATCTGGGACTACAAAAGAGATGCTCCAGCACATATATCTACTATTGGTGTTTGTTCTATGACTGACATAGCCAAGAAACCAACTGAAACGATTTGTGCACCACTCACTGTCTTTTTTGATGGTAGAGTTGATGGTCAAGTAGACTTATTTAGAAATGCCCGTAATGGTGTTCTTATTACAGAAGGTAGTGTTAAAGGTTTACAACCATCTGTAGGTCCCAAACAAGCTAGTCTTAATGGAGTCACATTAATTGGAGAAGCCGTAAAAACACAGTTCAATTATTATAAGAAAGTTGATGGTGTTGTCCAACAATTACCTGAAACTTACTTTACTCAGAGTAGAAATTTACAAGAATTTAAACCCAGGAGTCAAATGGAAATTGATTTCTTAGAATTAGCTATGGATGAATTCATTGAACGGTATAAATTAGAAGGCTATGCCTTCGAACATATCGTTTATGGAGATTTTAGTCATAGTCAGTTAGGTGGTTTAATCTTT

Nucleocapsid Sub-genomic RNA

GATTAAAGGTTTATACCTTCCCAGGTAACAAACCAACCAACTTTCGATCTCTTGTAGATCTGTTCTCTAAACGAACAAACTAAAATGTCTGATAATGGACCCCAAAATCAGCGAAATGCACCCCGCATTACGTTTGGTGGACCCTCAGATTCAACTGGCAGTAACCAGAATGGAGAACGCAGTGGGGCGCGATCAAAACAACGTCGGCCCCAAGGTTTACCCAATAATACTGCGTCTTGGTTCACCGCTCTCACTCAACATGGCAAGGAAGACCTTAAATTCCCTCGAGGACAAGGCGTTCCAATTAACACCAATAGCAGTCCAGATGACCAAATTGGCTACTACCGAAGAGCTACCAGACGAATTCGTGGTGGTGACGGTAAAATGAAAGATCTCAGTCCAAGATGGTATTTCTACTACCTAGGAACTGGGCCAGAAGCTGGACTTCCCTATGGTGCTAACAAAGACGGCATCATATGGGTTGCAACTGAGGGAGCCTTGAATACACCAAAAGATCACATTGGCACCCGCAATCCTGCTAACAATGCTGCAATCGTGCTACAACTTCCTCAAGGAACAACATTGCCAAAAGGCTTCTACGCAGAAGGGA GCAGAGGCGGCAGTCAAGCCTCTTCTCGTTCCTCATCACGTAGTCGCAACAGTTCAAGAAATTCAACTCCAGGCAGCAGTAGGGGAACTTCTCCTGCTAGAATGGCTGGCAATGGCGGTGATGCTGCTCTTGCTTTGCTGCTGCTTGACAGATTGAACCAGCTTGAGAGCAAAATGTCTGGTAAAGGCCAACAACAACAAGGCCAAACTGTCACTAAGAAATCTGCTGCTGAGGCTTCTAAGAAGCCTCGGCAAAAACGTACTGCCACTAAAGCATACAATGTAACACAAGCTTTCGGCAGACGTGGTCCAGAACAAACCCAAGGAAATTTTGGGGACCAGGAACTAATCAGACAAGGAACTGATTACAAACATTGGCCGCAAATTGCACAATTTGCCCCCAGCGCTTCAGCGTTCTTCGGAATGTCGCGCATTGGCATGGAAGTCACACCTTCGGGAACGTGGTTGACCTACACAGGTGCCATCAAATTGGATGACAAAGATCCAAATTTCAAAGATCAAGTCATTTTGCTGAATAAGCATATTGACGCATACAAAACATTCCCACCAACAGAGCCTAAAAAGGACAAAAAGAAGAAGGCTGATGAAACTCAAGCCTTACCGCAGAGACAGAAGAAACAGCAAACTGTGACTCTTCTTCCTGCTGCAGATTTGGATGATTTCTCCAAACAATTGCAACAATCCATGAGCAGTGCTGACTCAACTCAGG

387 Additional Nucleotides

CTAGAAGATCTCCTACAATATTCTCAGCTGCCATGGAAAATCGATGTTCTTCTTTTATTCTCTCAAGATTTTCAGGCTGTATATTAAAACTTATATTAAGAACTATGCTAACCACCTCATCAGGAACCGTTGTAGGTGGCGTGGGTTTTCTTGGCAATCGACTCTCATGAAAACTACGAGCTAAATATTCAATATGTTCCTCTTGACCAACTTTATTCTGCATTTTTTTTGAACGAGGTTTAGAGCAAGCTTCAGGAAACTGAGACAGGAATTTTATTAAAAATTTAAATTTTGAAGAAAGTTCAGGGTTAATAGCATCCATTTTTTGCTTTGCAAGTTCCTCAGCATTCTTAACAAAAGACGTCTCTTTTGACATGTTTAAAGTTT

1988 Additional Nucleotides

CTAGAAGATCTCCTACAATATTCTCAGCTGCCATGGAAAATCGATGTTCTTCTTTTATTCTCTCAAGATTTTCAGGCTGTATATTAAAACTTATATTAAGAACTATGCTAACCACCTCATCAGGAACCGTTGTAGGTGGCGTGGGTTTTCTTGGCAATCGACTCTCATGAAAACTACGAGCTAAATATTCAATATGTTCCTCTTGACCAACTTTATTCTGCATTTTTTTTGAACGAGGTTTAGAGCAAGCTTCAGGAAACTGAGACAGGAATTTTATTAAAAATTTAAATTTTGAAGAAAGTTCAGGGTTAATAGCATCCATTTTTTGCTTTGCAAGTTCCTCAGCATTCTTAACAAAAGACGTCTCTTTTGACATGTTTAAAGTTTAAACCTCCTGTGTGAAATTGTTATCCGCTCACAATTCCACACATTATACGAGCCGGAAGCATAAAGTGTAAAGCCTGGGGTGCCTAATGAGTGAGCTAACTCACATTAATTGCGTTGCGCTCACTGCCAATTGCTTTCCAGTCGGGAAACCTGTCGTGCCAGCTGCATTAATGAATCGGCCAACGCGCGGGGAGAGGCGGTTTGCGTATTGGGCGCTCTTCCGCTTCCTCGCTCACTGACTCGCTGCGCTCGGTCGTTCGGCTGCGGCGAGCGGTATCAGCTCACTCAAAGG CGGTAATACGGTTATCCACAGAATCAGGGGATAACGCAGGAAAGAACATGTGAGCAAAAGGCCAGCAAAAGGCCAGGAACCGTAAAAAGGCCGCGTTGCTGGCGTTTTTCCATAGGCTCCGCCCCCCTGACGAGCATCACAAAAATCGACGCTCAAGTCAGAGGTGGCGAAACCCGACAGGACTATAAAGATACCAGGCGTTTCCCCCTGGAAGCTCCCTCGTGCGCTCTCCTGTTCCGACCCTGCCGCTTACCGGATACCTGTCCGCCTTTCTCCCTTCGGGAAGCGTGGCGCTTTCTCATAGCTCACGCTGTAGGTATCTCAGTTCGGTGTAGGTCGTTCGCTCCAAGCTGGGCTGTGTGCACGAACCCCCCGTTCAGCCCGACCGCTGCGCCTTATCCGGTAACTATCGTCTTGAGTCCAACCCGGTAAGACACGACTTATCGCCACTGGCAGCAGCCACTGGTAACAGGATTAGCAGAGCGAGGTATGTAGGCGGTGCTACAGAGTTCTTGAAGTGGTGGCCTAACTACGGCTACACTAGAAGGACAGTATTTGGTATCTGCGCTCTGCTGAAGCCAGTTACCTTCGGAAAAAGAGTTGGTAGCTCTTGATCCGGCAAACAAACCACCGCTGGTAGCGGTGGTTTTTTTGTTTGCAAGCAGCAGATTACGCGCAGAAAAAAAGGATCTCAAGAAGATCCTTTGATCTTTTCTACGGGGTCTGACGCTCAGTGGAACGAAAACTCACGTTAAGGGATTTTGGTCATGAGATTATCAAAAAGGATCTTCACCTAGATCCTTTTAAATTAAAAATGAAGTTTTAAATCAATCTAAAGTATATATGAGTAAACTTGGTCTGACAGTTACCAATGCTTAATCAGTGAGGCACCTATCTCAGCGATCTGTCTATTTCGTTCATCCATAGTTGCCTGACTCCCCGTCGTGTAGATAACTACGATACGGGAGGGCTTACCATCTGGCCCCAGTGCTGCAATGATACCGCGAGACCCACGCTCACCGGCTCCAGATTTATCAGCAATAAACCAGCCAGCCGGAAGGGCCGAGCGCAGAAGTGGTCCTGCAACTTTATCCGCCTCCATCCAGTCTATTAATTGTTGCCGGGAAGCTAGAGTAAGTAGTTCGCCAGTTAATAGTTTGCGCAACGTTGTTGCCATTGCTACAGGCATCGTGGTGTCACGCTCGTCGTTTGGTATGGCTTCATTCAGCTCCGGTTCCCAACGATCAAGGCGAGTTACATGATCCCCCATGTTGTGCAAAAAAGCGGTTAGCTCCTTCGGTCCTCCGAT

---

**Supplemental Figure 1:**
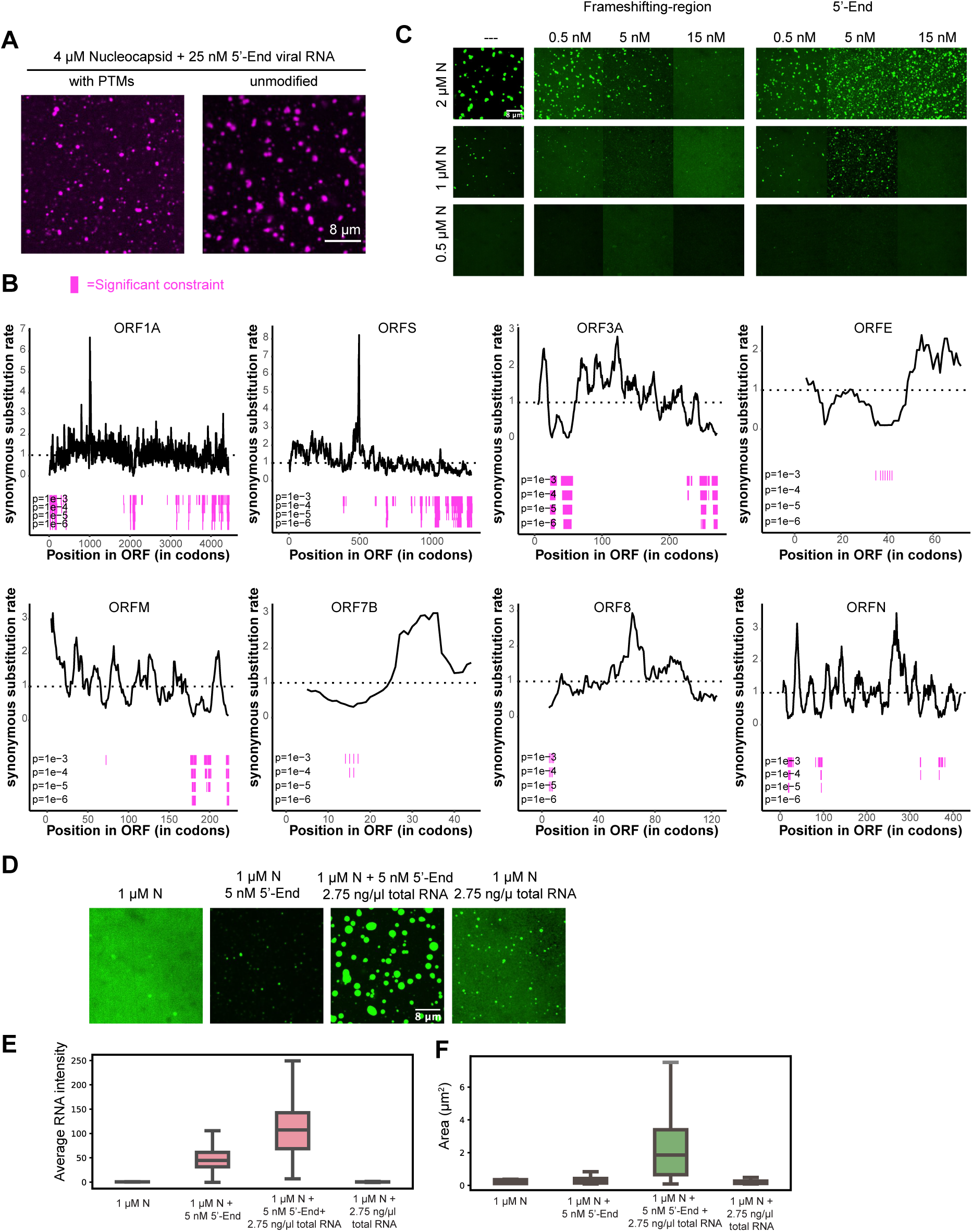
N protein LLPS is concentration and RNA species dependent. **A)** Unmodified N-protein and N-protein containing PTMs phase separate with viral RNA (cy3). **B)** Multiple regions across SARS-CoV2 genome (X axis) show elevated synonymous constraints. Y axis less than 1. Magenta bars indicate regions with significant synonymous constraints. Row 1 P<0.0001, Row 2 P< 0.00001, Row 3 P<0.00001, Row 4 P< 0.000001. **C)** Frameshifting RNA promotes dissolution of N-protein condensates (green) whereas 5’-End can drive phase separation relative to N-protein alone (left most panels). **D)** Non-viral lung cell RNA enhances LLPS with 5’-End. **E)** Quantification of average droplet RNA intensity from (D). **F)** Quantification of droplet area from (D). Scale bar, 8 µm unless otherwise noted

**Supplemental Figure 2:**
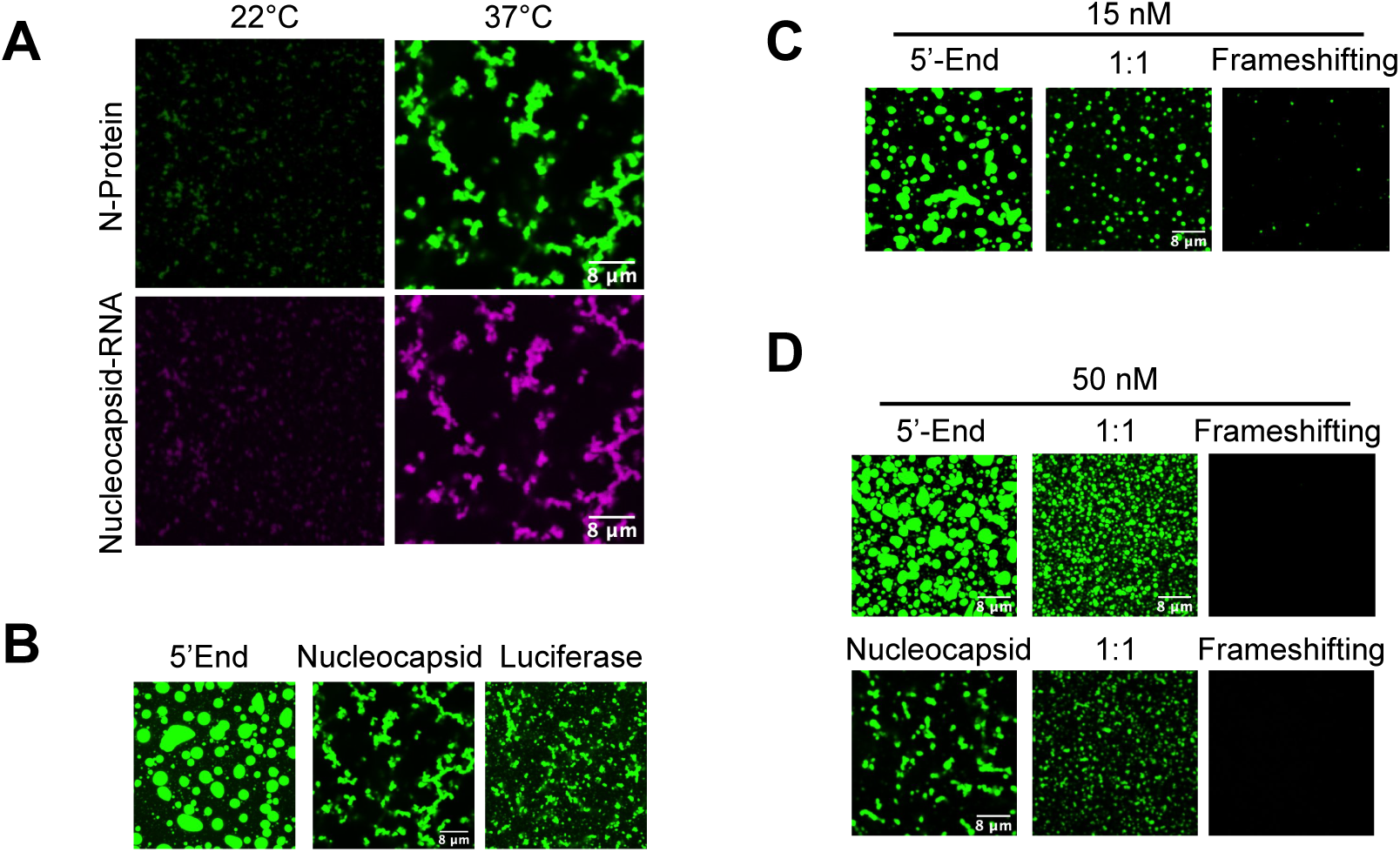
N protein LLPS is temperature dependent with RNA sequence encoded material properties. **A)** Nucleocapsid RNA phase separation with N-protein is temperature dependent. Nucleocapsid RNA (magenta) readily phase separates with purified N-protein (green) at 37°C, mammalian body temperature, but not at 22°C. **B)** 5’-End /N: protein (green) droplets are rounder than Nucleocapsid Luciferase RNA droplets. **C/D)** 5’-End or nucleocapsid RNA were mixed together with frameshifting RNA prior to N protein addition resulting in droplets of intermediate phenotypes (middle columns 1:1) compared to those of 5’-End (left) or frameshifting RNA alone (right). Scale bar, 8 µm unless otherwise noted.

**Supplemental Figure 3:**
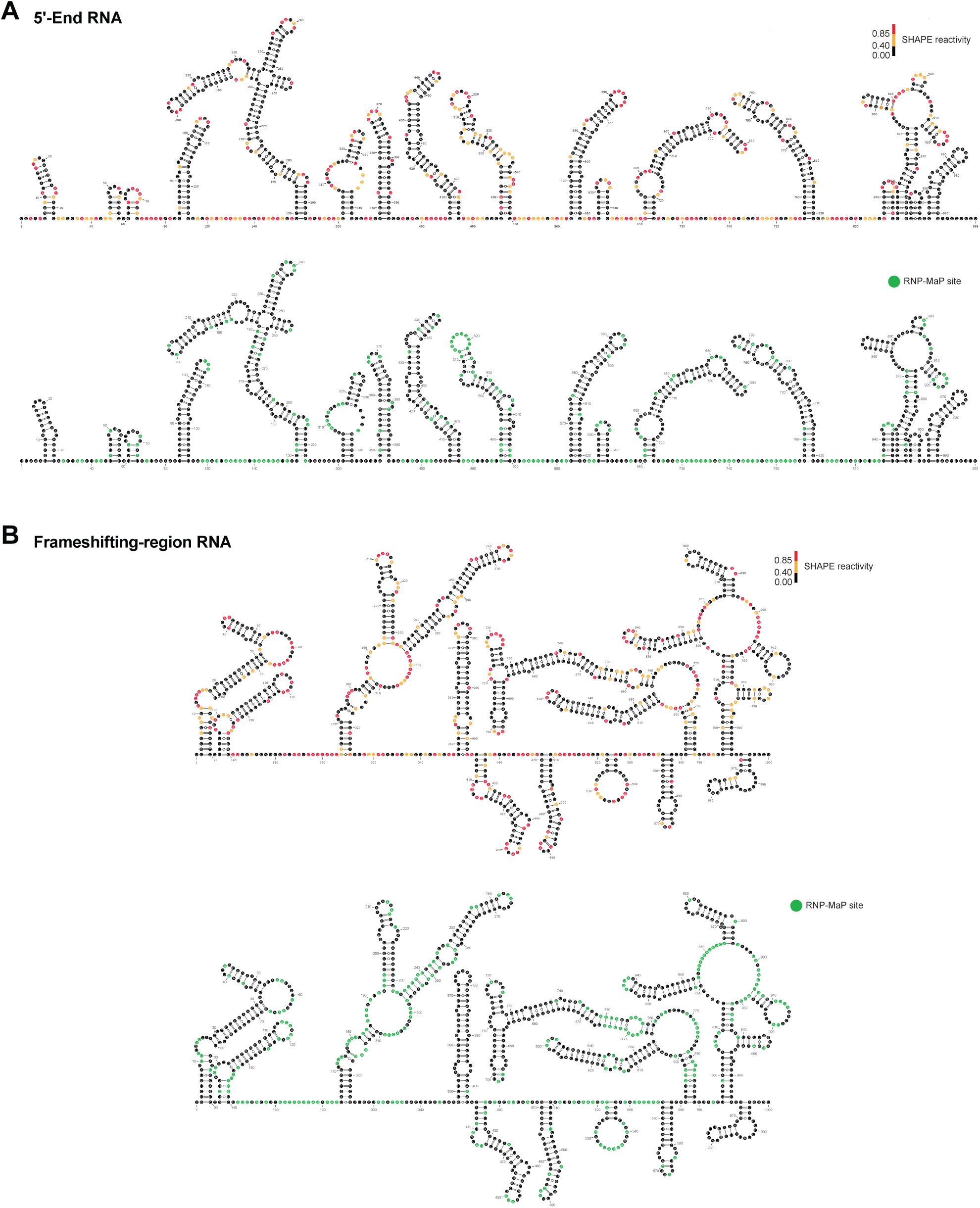
High-resolution RNA structures. SHAPE-directed secondary structure models for **(A)** the 5’-End RNA and **(B)** the Frameshifting-region RNA showing nucleotide identities and colored by SHAPE reactivity (upper panel, see scale) or RNP-MaP sites (lower panel). The lower panels of (A) and (B) are also shown in Fig. 3A and B.

**Supplemental Figure 4:**
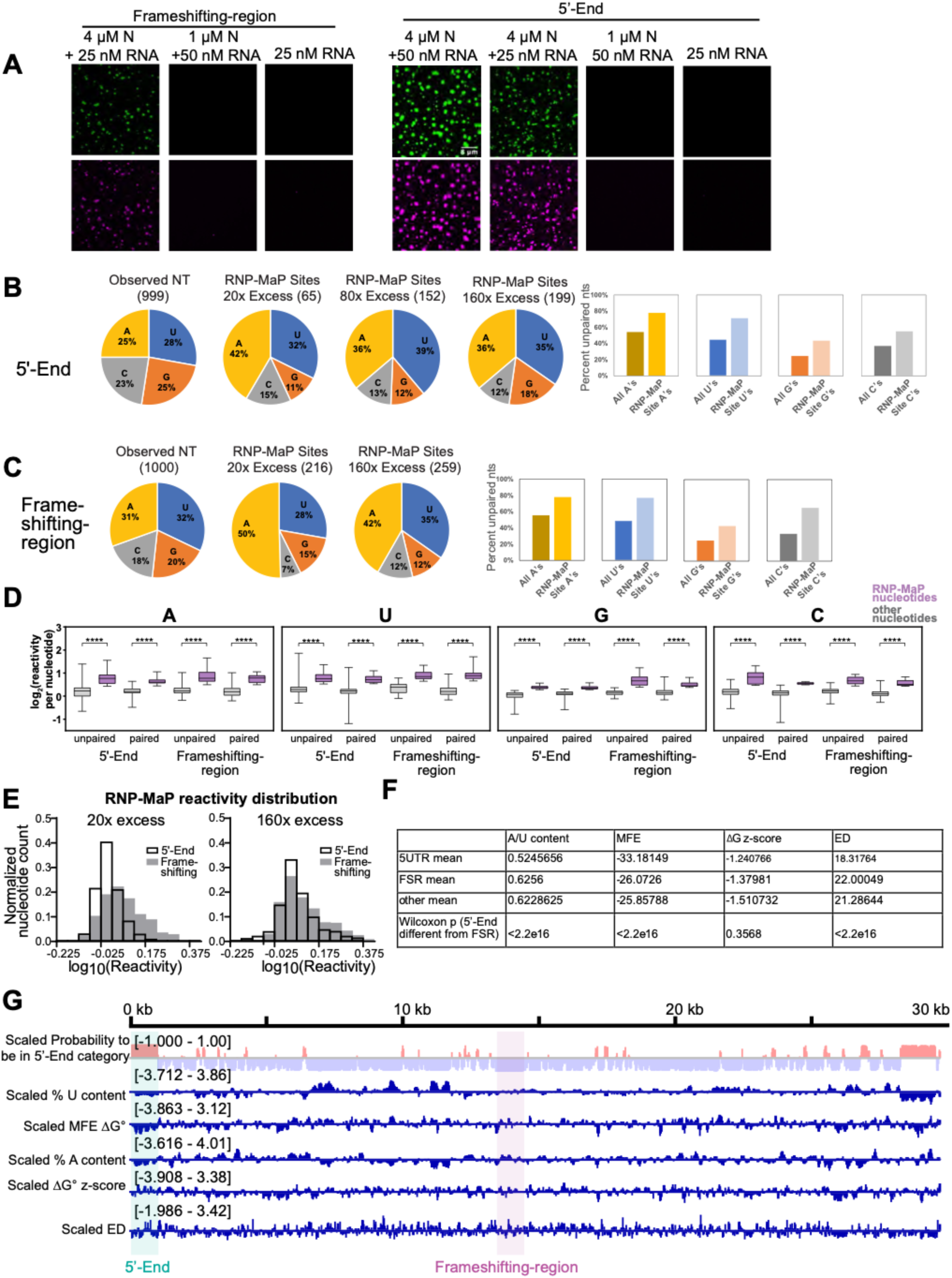
RNP-MaP reveals viral RNA-specific and condition-dependent N-protein binding site characteristics. **A)** Representative conditions for RNP-map experiments. Frameshifting-region and 5’-End RNAs (magenta) were mixed with N-protein (green) or droplet buffer and imaged prior to RNP-MaP experiments. Scale bar, 8 µm. **B)** N-protein–RNA interactions as defined by RNP-MaP sites are enriched in A/U nucleotides (pie charts) and unstructured nucleotides (bar graphs). Pie charts show RNP-MaP site distributions (in 160x conditions) as a function of nucleotide identity. Bar graphs show the percent of unpaired nucleotides in RNP-MaP sites compared to the full-length RNA. **C and D)** RNP-MaP detects N-protein–RNA interactions largely independent of nucleotide identity and local RNA structure. RNP-MaP site and non-site reactivities per nucleotide (A, U, G, C; in 160x conditions) were grouped as unpaired or paired and the reactivity per nucleotide was plotted. RNP-MaP sites (N-protein–RNA interactions) are significantly more reactive than non-sites (p < 1e-4, one-sided Kolmogorov-Smirnov test), while both unpaired and paired nucleotides have similar reactivities that suggest their unbiased detection (p > 0.06, one-sided Kolmogorov-Smirnov test). **E)** Comparison of RNP-MaP reactivity distributions for 5’-End and Frameshifting-region RNAs in both diffuse (20x excess) and condensed (160x) conditions reveal the Frameshifting-region RNA to be more broadly interacting with N-protein (larger distribution of higher reactivity nucleotides) in both conditions. **F)** RNA features used to classify 5’-End-like and Frameshifting-region-like regions in the gRNA. **G)** Genome similarity to 5’-End and Frameshifting-region. Mean for each feature is computed over all 120 base pair windows with center in the region of interest. MFE, dG z-score, and ensemble diversity are defined in (18).

**Supplemental Figure 5:**
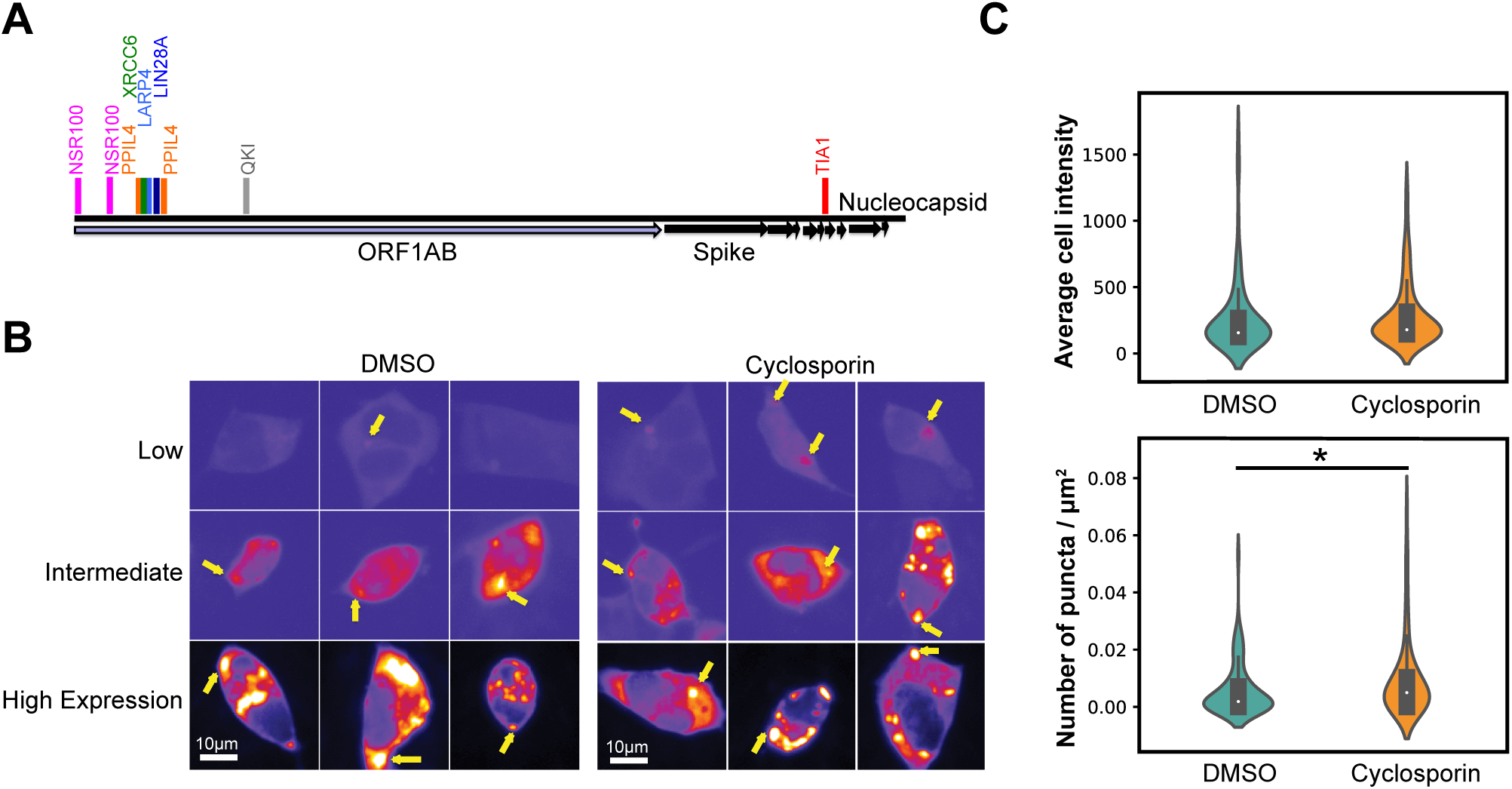
Cyclosporin treatment increase N-protein puncta relative to vehicle. **A)** Approximate locations for Seqweaver predictions for RNA-binding proteins interaction with the SARS-CoV2 genomic RNA. **B)** Relative to vehicle, cyclosporin treatment induces puncta (yellow arrows) in cells with lower N:protein (Fire LUT) expression. Scale bar is 10 µm. **C)** N protein whole cell intensity is equivalent, but number of puncta/ square micron increases >1.6X with cyclosporin treatment. * p<0.05.

## Author contributions

C.A.R. and C.I. contributed equally to this work and order of authorship was determined alphabetically. C.A.R., C.I. and A.S.G. conceptualized the project, designed experiments, prepared figures, drafted and edited the manuscript. C.A.R. and C.I. also performed experiments and analyzed data; M.A.B. designed and performed experiments and computational analyses, analyzed data, prepared figures, and edited the manuscript. K.M.W. designed experiments and analyzed data, and edited the manuscript; G.A.M. designed and performed experiments, analyzed data, edited the manuscript and performed computational analyses; R.S. designed and performed computational analysis and edited the manuscript; I.J. supported computational analyses; C.T. provided fruitful discussion and edited the manuscript; O. G. T. provided support for R. S. and C. T.; I.R. and C.P. designed and performed computational analysis; A.B. performed fruitful literature research on cyclosporine. E.F., Y.J.H., T.P.S. and R.B. harvested viral RNA.

## Data and materials availability

All data are available upon request from C.A.R., C.I. or A.S.G.

## Acknowledgements

We thank Rick Young, Phil Sharp, Alex Holehouse, Kathleen Hall, Andrea Sorrano, Ahmet Yildez and their lab members for sharing data and discussions, David Adalsteinsson for his help with ImageTank software, Ian Seim for analysis consultation and discussions, Benjamin Stormo for critical reading of the manuscript, Alain Laederach for initial discussion on genomic sequence, Chase Weidmann for initial discussions planning RNP-MaP experiments, and James Iserman for essential logistical support. A.S.G., C.I. and C. A. R. were supported by NIH R01GM081506 and an HHMI faculty Scholar Award, C.A.R. was supported by NIH T32 CA 9156-43, F32GM136164 and L’OREAL USA for Women in Science Fellowship. The work by RS, CYP, ASB, and CLT is supported by NIH grants R01HG005998, U54HL117798 and R01GM071966, HHS grant HHSN272201000054C and Simons Foundation grant 395506 to O.G.T. K.M.W. and M.A.B. were supported by US National Institutes of Health (R35 GM122532 to K.M.W.). M.A.B. was supported by a Ruth L. Kirschstein Postdoctoral Fellowship (F32 GM128330).

